# Deconstructing empirical fitness seascapes across scales of granularity

**DOI:** 10.64898/2026.02.04.703871

**Authors:** Swathi Nachiar Manivannan, C. Brandon Ogbunugafor

## Abstract

The fitness landscape metaphor remains resonant in evolutionary theory and has facilitated the birth of newer concepts—like the fitness seascape—that consider the role of environmental context in shaping the dynamics of evolution. Since the emergence of the fitness seascape, it has appeared in several studies that examine how different and fluctuating environments shape evolutionary outcomes. Despite a growing interest in these topics, we lack comprehensive examinations of the role of environmental context in shaping features of fitness seascapes. In this study, we address this gap by deconstructing empirical fitness seascapes across scales of granularity: mutational steps, loci, locus interactions, alleles, trajectories, and entire seascapes. For each, we examine how environmental context influences qualitative and quantitative aspects of seascapes, and find that they change appreciably, with patterns that are specific to individual systems of study. In summary, we reflect on the implications of the seascape metaphor with respect to the incorporation of environmental effects into theoretical population genetics, for understanding how the environment shapes evolution in disease systems, and for contemporary bioengineering excursions.

## 1 Introduction

The fitness or adaptive landscape is a central metaphor in evolutionary theory that likens the relationship between genotypes and phenotypes to a physical surface. It has been an integral part of evolutionary genetics since its formal introduction by Sewall Wright in 1932 (*1*), and has spawned a subfield focused on how the shape (or topography) of fitness landscapes dictates the dynamics of evolution at the molecular scale (*2*–*7*). It has also helped to develop detailed pictures of phenomena such as adaptive trajectories, epistasis, evolvability, and various other forces operating on molecular evolution (*8*–*12*). The strength of the fitness landscape metaphor lies in its relative simplicity: despite the high-dimensionality of genotype-phenotype space, most canonical studies of fitness landscapes have focused on static environments.

However, for many years, a body of work has emerged that examines the case where the fitness landscape topography changes across environments. The ambition here is to transform the study of fitness landscapes into a more realistic instrument, as the living world can be safely described as a highly capricious, multi-environment setting. This notion was first formalized by David Merrell as the “fitness seascape,” an iteration of the fitness landscape that considers how environmental context shapes evolution at the molecular level (*13*). Ville Mustonen and Michael Lässig then introduced a formal framework for the study of adaptation as a non-equilibrium phenomenon in fluctuating environments (*14*) to reflect the biological reality of adaptive evolution as well as in the context of anthropogenically driven environmental changes (*15*). This has culminated in a literature that has explored how different environmental contexts craft adaptive trajectories and landscape topographies to answer questions about adaptive evolution (*16*–*23*), niche construction (*24,25*), and speciation (*26,27*). However, evolution in dynamic environmental contexts remains relatively understudied, which has amplified the need for a better understanding of how contextual changes can drive adaptive evolution (*28*–*31*).

The richness of the fitness landscape metaphor resides in the number of different questions one can ask of it. For example, previous studies have used empirical fitness landscapes to examine questions as disparate as the distribution of fitness effects (*32,33*) to speciation (*34*). This breadth of inquiry is possible because fitness landscapes are intrinsically composed of different kinds of biological information, which are relevant to different areas of evolutionary and population genetics. Studies have used fitness landscapes to ask questions about mutational steps (*35,36*), plasticity (*37*), epistasis (*15,33,38*), and landscape topography (*39*). In this study, we propose a framework for computing how changing environmental contexts influence different mechanistic features of seascapes across scales of granularity. Specifically, we examine the manner in which the environmental context influences mutational steps, mutation effects, mutation interactions (epistasis), alleles, trajectories, and descriptions of entire landscapes, such as ruggedness. In doing so, we attempt to deconstruct the anatomy of molecular evolution.

Our framework comes at a time when evolutionary biologists are acquiring more and more empirical data and beginning to uncover the intricacies of how genetic information maps onto phenotypic traits and the fitness of organisms. We first discuss how our framework deconstructs evolvability and related features at different scales of the fitness landscape before testing our framework on an assortment of empirical fitness seascapes. Finally, we communicate the implications of our findings and the relevance of our framework to broader discussions in evolutionary genetics and evolvability.

## 2 Materials and Methods

### 2.1 Fitness seascapes data

A majority of the empirical fitness landscapes used in this study were consolidated and analyzed in a 2018 study. (*40*). These landscapes were also supplemented by others focused on dihydrofolate reductase (DHFR) (*41*– *43*) and a TEM-ω-lactamase in different bacterial species (*44*). Table 1 summarizes the 10 fitness seascapes used in this study and includes information on the study from which the seascape dataset came, the description of what was measured as a fitness proxy, the number and type of environmental and/or phenotypic contexts in the seascape dataset, and the number of loci in each seascape. Several seascape datasets include logtransformed or normalized versions of phenotypic traits to facilitate interpretation, since most metrics in our framework assume that measurements of fitness are on an additive scale. These datasets provide a diverse and representative foundation for demonstrating how our framework can be applied to seascapes in general. For the purposes of this paper, we provide a detailed discussion of two datasets: the Long-Term Evolution Experiment (LTEE) seascape, which consists of 32 genotypes of *Escherichia coli* generated from the first five beneficial mutations fixed in the LTEE grown in three different conditions (*45,46*), and the *Plasmodium falciparum* DHFR pyrimethamine seascape, which consists of 16 genotypes of *P. falciparum* generated from four important drug-resistant mutations grown in a concentration gradient of the antimalarial drug pyrimethamine (*43,47*). This choice is intentional, as they represent fitness seascapes from a diversity of organisms, though both are microbial. However, more importantly, they demonstrate two different ways in which seascapes can be constructed: The DHFR pyrimethamine seascape is continuous and quantitative, while the LTEE seascape is categorical. This demonstrates flexibility in the use cases of the methods we have provided and the greater fitness seascape idea. In addition, we also provide a discussion of two other datasets (*44,48*) in the supplementary information.

**Table 1.** Table of fitness seascapes used in this study. Datasets that are discussed in the main section of the paper are bolded, datasets that are discussed in the supplementary information are bolded and italicized.

| **Dataset** | **Description** | **Number of environments/phenotypes** | **Loci** |
| --- | --- | --- | --- |
| Human immunodeficiency virus replication (49) | $\log_{10}$(replicative capacity of HIV in host cells) | 2 (CCR5+ cells, CXCR4+ cells) | 5 |
| DHFR proteostasis (42) | $\ln$(Protein abundance) and $\ln(IC_{50})$ of DHFR variants across different bacterial species and proteostatic backgrounds | 18 (2 phenotypes ($IC_{50}$ & DHFR protein abundance) × 3 bacterial species (*Escherichia coli*, *Listeria grayi*, *Chlamydia muridarum*) × 3 proteostatic backgrounds (WT, *groEL*<sup>+</sup>, $\Delta lon$)) | 3 |
| *Saccharomyces cerevisiae* growth rates (50) | $\ln$(Growth rate of *S. cerevisiae*) | 2 (haploid, diploid) | 6 |
| **LTEE (45,46)** | $\ln$(Relative fitness of *E. coli*) | 3 (DM25, DM25 + EGTA, DM25 + guanazole) | 5 |
| *bla*<sub>TEM</sub> **cefotaxime resistance (44)** | Relative resistance level of *bla*<sub>TEM</sub> alleles to cefotaxime in different bacterial species | 3 (*E. coli*, *Salmonella enterica*, *Klebsiella pneumoniae*) | 5 |
| DHFR pyrimethamine resistance in *Plasmodium* spp. (51,52) | $\ln$(Pyrimethamine $IC_{50}$ in *Plasmodium* spp.) | 3 (*P. falciparum* assayed in *E. coli*, *P. vivax* assayed in *E. coli*, *P. vivax* assayed in *S. cerevisiae*) | 4 |
| **β-lactam resistance (48)** | $\ln$(Growth rate of *E. coli* in different β-lactams) | 7 (amoxicillin, amoxicillin/clavulanic acid, cefotaxime, cefotetan, cefprozil, ceftazidime, piperacillin/tazobactam) | 4 |
| **DHFR resistance in pyrimethamine / cycloguanil (43,47)** | $\ln$(Growth rate of *Plasmodium falciparum*) in different concentrations of pyrimethamine or cycloguanil | 20 (10 different concentrations of pyrimethamine and cycloguanil each) | 4 |
| Sesquiterpene synthase biochemistry (53) | % production of different products by sesquiterpene synthase mutants in *Nicotiana tabacum* | 4 (premnaspirodiene, 4-epi-eremophilene, 5-epi-aristolochene, minor products) | 6 |
| *bla*<sub>TEM</sub> cefotaxime / piperacillin resistance (54,55) | $\log_{\sqrt{2}}$(mean MIC) of *bla*<sub>TEM</sub> alleles in *E. coli* | 3 (cefotaxime, piperacillin, piperacillin + clavulanic acid) | 5 |

### 2.2 Framework for deconstructing fitness seascapes

Figure 1 provides a graphical overview of our proposed framework for studying features of fitness seascapes and how they vary across different contexts. Here, we discuss how to compute metrics of different features of fitness seascapes, starting from the scale of mutational steps, then progressing to loci, alleles, and seascape-wide features.

**Fig. 1.**
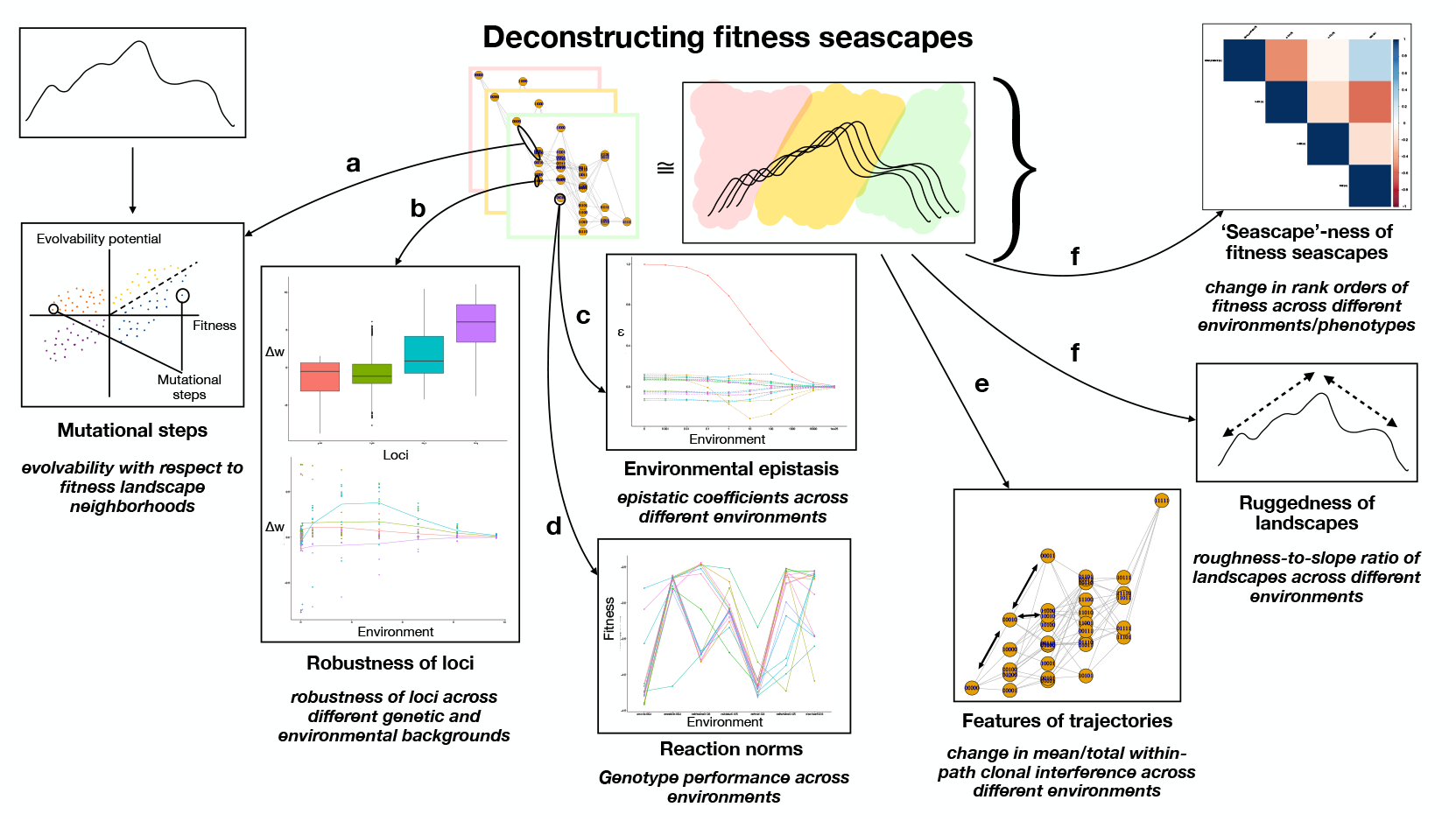
A framework for deconstructing the study of fitness seascapes across different scales of granularity (biological organisation). From left to right: (a) measuring evolvability potential of mutational steps, (b) genetic and environmental robustness of loci, (c) environmental epistasis at the scale of interactions between loci, (d) reaction norms at the scale of alleles, (e) the shape of accessible trajectories and the speed of evolution across those trajectories, and ruggedness and ‘seascape’-ness to measure topography at the seascape scale.

We note that the hierarchy that we have imposed is neither essential nor exhaustive. Empirical fitness landscapes/seascapes are not objectively organized into scales based on granularity. Our presentation is largely for organizational purposes. Relatedly, while we have chosen scales and methods for which there are published methods, our organization does not represent every possible sort of analysis applied to a fitness seascape^1^. We urge those interested to apply other analysis methods to understand how environments shape fitness seascape features.

#### 2.2.1 Mutational steps: evolvability potential

At the level of mutational steps, we can consider how the relationship between fitness and evolvability varies across different environmental contexts. We adapt the “evolvability-enhancing mutations” metric proposed in Wagner (2023) (*36,56*) to consider how accessibility to mutational neighborhoods with a higher mean fitness varies across contexts (Figure S1).

To measure evolvability and fitness at the mutational step scale, we first break down each fitness seascape to their individual landscapes (i.e., treating every unique environmental/phenotypic context as an individual landscape). For each landscape, we consider every possible pair of genotypes that are 1 mutational step away from each other (e.g. 0000–0001) and arbitrarily assign one genotype to be the wild-type (wt), and the other to be the mutant (m). We consider both directions of the mutational step separately (i.e., A → B and B → A are considered as two different steps, but will have opposite magnitudes).

The fitness effect of the mutational step is Δ*w* = *w* (*m*) −*w* (*wt*), where *w* (*wt*) is the fitness of the wt genotype and *w* (*m*) the fitness of the m genotype. It then follows that beneficial mutational steps have Δ*w* > 0 and deleterious mutational steps have Δ*w* < 0.

The evolvability potential of the mutational step is calculated by comparing the mean fitness of the 1-mutational step neighborhood of the m genotype to the 1-mutational step neighborhood of the wt genotype, i.e.,

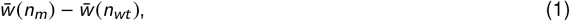

where 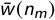 is the mean fitness of all genotypes that are a 1-mutation neighbor of the m genotype *excluding the wt genotype*, and 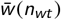 is the mean fitness of all genotypes that are a 1-mutation neighbor of the wt genotype *excluding the m genotype*. To categorize mutational steps as evolvability-enhancing (EE)/non-evolvability-enhancing (nonEE), we rely on the following inequality:

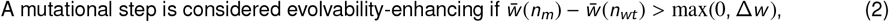

since beneficial mutational steps would trivially meet the condition of 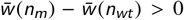. Following which, for every individual fitness landscape within and across seascapes, we plot and color-code all mutational steps based on whether they are beneficial/deleterious and EE/nonEE (Figure S1), thus generating mutational step profiles for every seascape. Note that this calculation applies to quantitative properties of mutational steps, independent of what locus undergoes mutation.

#### 2.2.2 Loci: genetic and environmental robustness

Next, we consider how features of genetic loci vary across environmental contexts, specifically focusing on the genetic and environmental robustness of loci. Adapting from a prior study of reverse evolution in *Plasmodium vivax* DHFR (*57*), we calculate genetic and environmental robustness as such:

We first calculate the fitness effect of a mutation, *ϵ*, by taking the mean difference between the fitness of all pairs of allele *j* ***not*** carrying ϵ and the one-step neighbor carrying mutation *ϵ*,

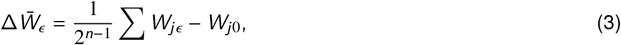

where *ϵ* corresponds to the mutant allele in a given locus and *n* denotes the number of loci in the seascape. For example, in a 4-locus system, *ϵ* could refer to one of the following: ‘1***’, ‘*1**’, ‘**1*’ or ‘***1’. Genetic robustness is then defined as 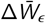 summarized for a locus across all genetic backgrounds ***and*** environmental contexts, whereas environmental robustness is determined by comparing how 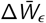 computed for a locus across all genetic backgrounds varies between environmental contexts.

#### 2.2.3 Interactions between loci: Environmental epistasis and the mutation-effect reaction norm (MuRNs)

Building on individual loci, we can then explore how the interactions between loci (i.e., mutation effects and epistasis (G × G interactions)) shift across contexts. The concept of “environmental epistasis” or environmentally-dependent epistatic interactions have been observed across a number of empirical systems, where the effect of a single mutation, or the magnitude and sign of epistatic interactions between multiple mutations can vary depending on the context in which the system exists (i.e., G × E and G × G × E interactions) (*48,58,59*). A tractable way of capturing these interactions (and therefore how variable seascapes are across environments) is through a “mutation-effect reaction norm” (MuRN) (*43*). Computing the MuRN for a seascape involves the Walsh-Hadamard transform to compute coefficients corresponding to the sign and magnitude of interactions between mutations (*38,40*). We refer readers to various studies on the methods and how they have been used to measure higher-order epistasis (*38,60*) for a more detailed discussion, but we provide a brief summary of the methodology used for computing the MuRN below:

For every environmental context in a given seascape, we compute the single-mutation effect and epistatic interaction coefficients as follows:

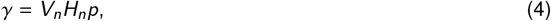

where *γ* = the Walsh coefficient vector, *n* = number of loci in the fitness seascape, *V*_*n*_ = a normalizing diagonal matrix for 2^*n*^ genotypes,*H*_*n*_ = Hadamard matrix for 2^*n*^ genotypes, and *p* = the vector of fitness values arranged numerically based on genotypes depicted in binary notation. *V*_*n*_ and *H*_*n*_ are defined recursively as such:

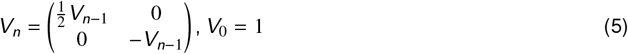

and

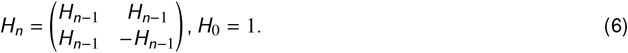

For example, in a 3-locus landscape, the phenotypes in *p* will be arranged in the following manner (corresponding to the respective genotypes): [000, 001, 010, 011, 100, 101, 110, 111]. In the MuRN, we can consider not only the coefficients of the individual interactions, but also the mean interaction coefficients within an order to allow for comparison of interactions between orders. This is done by computing the absolute mean coefficient for every order of interaction,

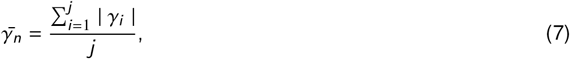

where *n* = the order of interaction, and *j* = the number of epistatic coefficients that represent the *n*th order of interaction. We note that these methods were pioneered for use on fitness landscapes by Daniel Weinreich and colleagues (*38*), and similar methods have been utilized in other studies of genotype-phenotype maps (*60*–*62*).

#### 2.2.4 Alleles: Reaction norms

At the allele scale, we can compute reaction norms to describe how the performance (phenotype or fitness) of genotypes is shaped by the environment (*63*). In the scope of this study, we do a qualitative comparison of how each genotype performs with respect to other genotypes in different environments; however, we note that more quantitative treatments of reaction norms can be and have been utilized in quantitative genetics and phenotypic plasticity, among other fields (*58,64*–*66*). Furthermore, we also take a more quantitative analysis of how the reaction norms manifest at the landscape scales, where we compare how the rank orders of genotypes change across environments (see discussion on landscape topography).

#### 2.2.5 Features of trajectories: Within-path competition and the speed of evolution

Having examined properties of mutational steps, loci, and alleles, we now look at features at a higher scale of granularity: the evolutionary trajectory. Trajectories are observational summaries of individual mutational steps that occur in statistical space. Nonetheless, molecular evolution does occur along some evolutionary trajectories (or “paths” or “adaptive walks”) more readily than others, and so they are a meaningful scale to examine.

We can consider how adaptive evolutionary trajectories can vary across different contexts as a consequence of topographical changes. We adapt the within-path competition metric, *C*_*w*_, proposed in a previous study as a proxy for the speed of adaptive evolution of antimicrobial resistance across landscapes (*67*). The formula for computing *C*_*w*_ is as follows:

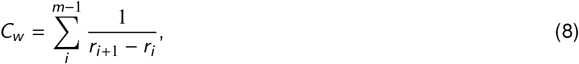

where *r*_*i*_ = growth rate (or any other fitness proxy) of genotype *i* and *m* = number of genotypes in a given pathway. We only compute *C*_*w*_ for all possible upward trajectories ^2^.

To compare how the speed of evolution (as measured by within-path competition) varies across different contexts, we generalize *C*_*w*_ across the entire landscape, accounting for every possible beneficial trajectory in the landscape, by calculating *C*_*w*(tot)_ and *C*_*w*(avg)_, where

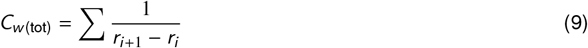

for all beneficial mutational steps across the landscape, and

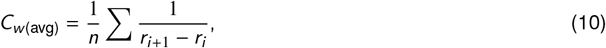

where *n* = number of beneficial mutational steps across the landscapes.

#### 2.2.6 Topographical features

Finally, at the landscape scale, we can consider how topographical features of seascapes vary across contexts.

##### Topography: ‘seascape-ness’ of fitness seascapes

Given the support for the role of environmental context in shaping seascape features at finer scales (e.g., mutational steps, alleles), we now ask how seascape topography changes in different contexts. One way in which we can quantify how much fitness landscape topography fluctuates is to do a pairwise comparison of the rank orders of genotypes in two different contexts, using Kendall’s τ rank correlation coefficient. Kendall’s τ ranges from -1 to 1, where 1 indicates a perfect agreement of rank orders and -1 a perfect disagreement of rank orders.

##### Topography: ruggedness of fitness seascapes

Another way of measuring the topography of fitness landscapes is to quantify how rugged a landscape is. We can extend this to fitness seascapes by measuring how the ruggedness of landscapes changes across different contexts, using the roughness-to-slope ratio for measuring ruggedness (*39,68*). This is done by fitting a multi-dimensional linear model of fitness to loci:

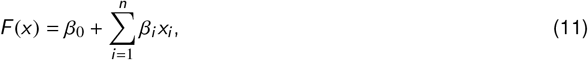

where *F* (*x*) = fitness, *x*_*i*_ = state at locus *i*, *n* = number of loci. The roughness is the mean squared error of the regression model,

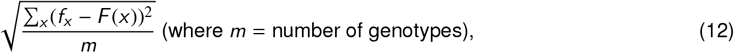

while the slope is the mean of absolute values of linear coefficients,

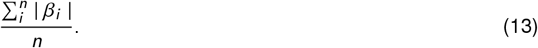

## 3 Results

In this section, we discuss how the various properties discussed in our framework vary within and across seascapes. At each scale, we illustrate this by primarily discussing the DHFR pyrimethamine (*43,47*) and LTEE (*45,46*) seascapes, and also discuss two other seascapes in the Supplementary Information (*44,48*). Finally, we briefly discuss other approaches for studying fitness seascapes, that were not mentioned in our framework but could be integrated nonetheless (also summarized in Table 2).

**Table 2.** A blueprint for studying features of fitness seascapes, discussing in detail the framework illustrated in Figure 1. Here, we discuss the scales of deconstruction, specific metrics measured at every scale, and literature that formally introduced these metrics and related concepts.

| Scale | Description of metric | References |
| --- | --- | --- |
| (a) Mutational step | **Fitness effect** ($w(m) - w(wt)$) and **evolvability potential** ($\bar{w}(n_m) - \bar{w}(n_{wt})$) of individual mutational steps to quantify accessibility to mutational neighbourhoods with higher mean fitness | (36) |
| (b) Locus | **Robustness** ($\Delta \bar{w}$) of loci across genetic and/or environmental backgrounds | (57) |
| (c) Locus interaction | **Consideration of environmental epistasis**, measured by:<br>• **Mutation-effect reaction norm (MuRN)** to understand how interactions between loci change across environmental contexts<br>• **Global epistasis × environment interactions** to predict the fitness effect of mutations based on genomic and environmental context | (43,58,59,75,76) |
| (d) Allele | **Reaction norm** to understand how the phenotypic trait values/fitness of individual genotypes vary across different contexts | (43,63,64) |
| (e) Trajectory | **Within-path competition** ($C_w$) to quantify how adaptive trajectories and the speed of adaptive evolution can vary across different environmental contexts | (67) |
| (f) Landscape topography | • **'Seascape'-ness** by comparing rank orders of genotypes between any two different environments<br>• **Roughness-to-slope ratio** to measure the ruggedness of landscapes and how they vary across environments<br>• **Edge-flip fraction** to quantify topographical shifts between any two different environments | (39,68,71,77) |
| Other theoretical and empirical measures | • **Fixation probability of novel mutations** in fluctuating environments<br>• Evolution of **mutation rates/mutator phenotypes** in fluctuating environments<br>• **Fitness variance and 'environmental memory'** of mutant strains when evolving in fluctuating environments<br>• **Distribution of fitness effects, distribution of diversity (clonal size distributions)** across different environments | (16,78–80) |

### 3.1 Properties of mutational steps

Across all the data sets that we studied, we observe that each seascape has unique mutational step profiles, both in terms of the proportions of different types of mutational steps, as well as the shapes of the mutational step profiles, as measured through correlation between evolvability potential and fitness effect. Notably, for any given seascape, these profiles can vary across environments, suggesting that accessibility to mutational neighborhoods of greater fitness can vary with context (Figure 2), which has implications for evolvability (*36*).

**Fig. 2.**
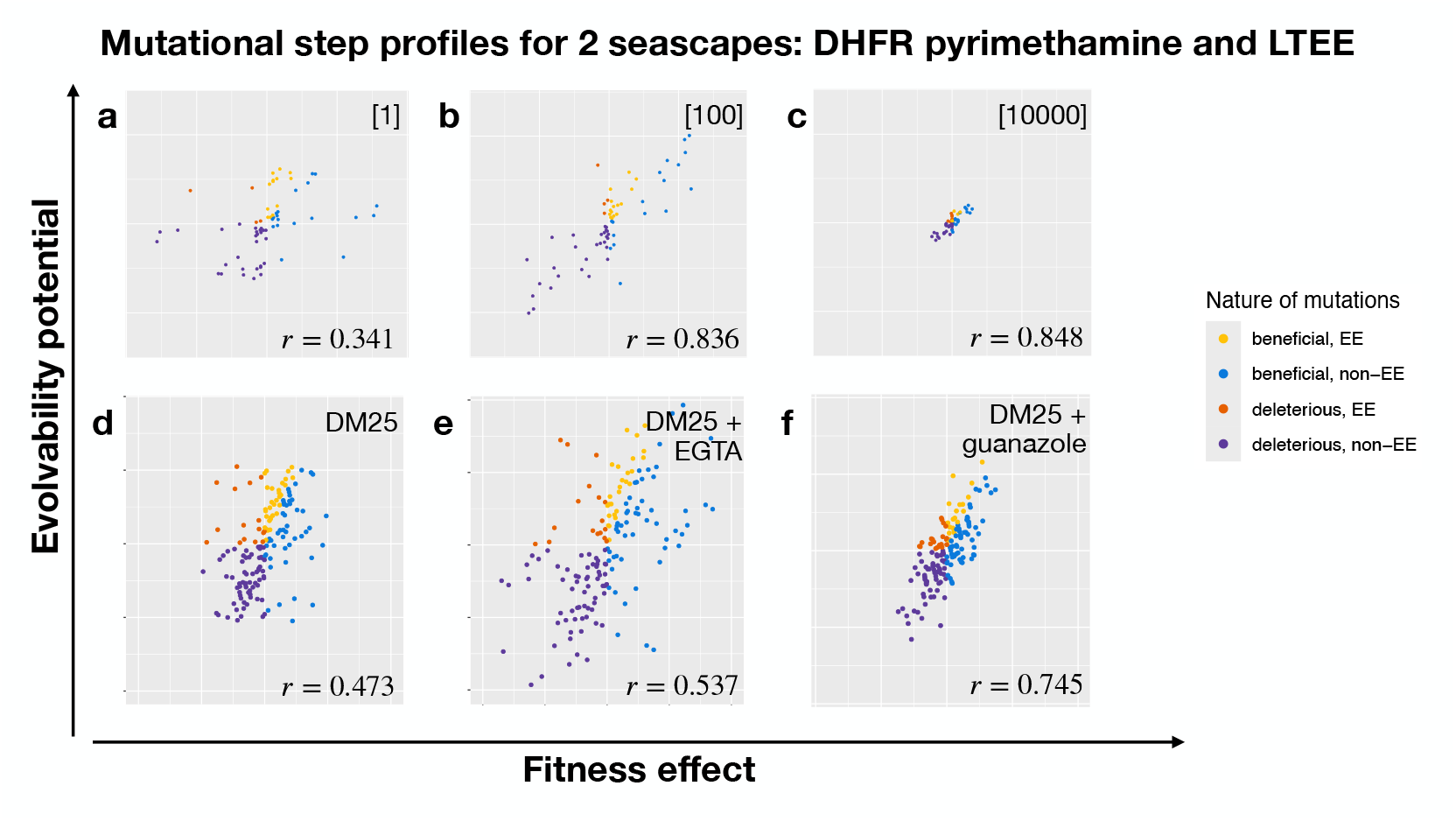
Select mutational step profiles from the DHFR pyrimethamine and LTEE seascapes. The DHFR pyrimethamine seascape maps the growth rates of 2^4^ DHFR genotypes in *P. falciparum* across a concentration gradient of pyrimethamine, whereas the LTEE seascape maps the growth rates of 2^5^ genotypes in *E. coli* (generated from the first five beneficial mutations of the Long-Term Evolution Experiment) across 3 different growth conditions. Mutational steps are color-coded as such: yellow - beneficial, EE; orange - deleterious, EE; purple - deleterious, nonEE; blue - beneficial, nonEE. a-c: DHFR pyrimethamine seascape, mutational step profiles for pyrimethamine concentrations of (a) 1, (b) 100, and (c) 10000 *µ*g ml^−1^. Subfigures a-c are plotted on x and y-axes with the same scale. d-f: LTEE seascape, mutational step profiles for (d) DM25, (e) DM25 + EGTA, and (f) DM25 + guanazole. For each profile, the Pearson’s correlation coefficient (*r*) between fitness effect and evolvability potential is also noted in the bottom right corner. Subfigures d-f are plotted on x and y-axes with the same scale.

### 3.2 Properties of loci

At the locus scale, we observe that genetic and environmental robustness manifest differently across different seascapes. For example, in the DHFR pyrimethamine seascape, we observe that the fourth locus (I164L) has a lower 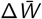 than all the other loci (1-way ANOVA followed by Tukey’s post-hoc test, Figure 3a). However, when considering 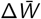 in particular environmental contexts, a two-way ANOVA (fitness ∼ locus * environment) suggests that the third locus (S108N) has a higher 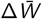 in pyrimethamine concentrations of 10*µ*g ml^−1^ (*p* = 0.00521), 100*µ*g ml^−1^ (*p* = 0.00180), and 1000*µ*g ml^−1^ (*p* = 0.0240) (Figure 3b). In the LTEE seascape, we observe that all five loci have similar 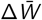 averaged across all genetic backgrounds and environments (1-way ANOVA followed by Tukey’s post-hoc test, Figure 3b), but a two-way ANOVA (*p* = 0.0002182) suggests that loci 1 (*rbs*) (*p* = 0.00495) and 4 (*glnUS*) (*p* = 0.00986) have higher 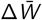 tance seascape, all four loci have similar 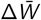 values than loci 2, 3 and 5 in DM25 + EGTA. In the *β*-lactam resisaveraged across all genetic backgrounds and environments (1-way ANOVA, *p* = 0.305), but when we look at 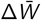 of the loci for each environment, the mutation G238S in the 3rd locus has a higher 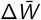 in certain environments, as demonstrated through a two-way ANOVA that examined the effect of locus and environment on 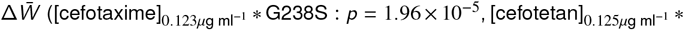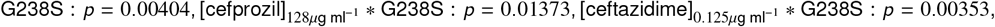 Figure S2a and b). On the other hand, in the bla_*TEM*_ cefotaxime resistance seascape, we observe that of the 5 loci, the mutation G238S in the 5th locus has the highest 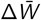, followed by the mutation E104K in the 3rd locus; these differences in 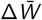 compared to the other loci are statistically significant (1-way ANOVA followed by Tukey’s post-hoc test, Figure S2c) and this holds across the three different species environments (*Escherichia coli, Klebsiella pneumoniae, Salmonella enterica*) (2-way ANOVA: locus, *p* < 2.2 10^−16^; species, *p* = 0.412; locus*species, *p* = 0.966, Figure S2d).

**Fig. 3.**
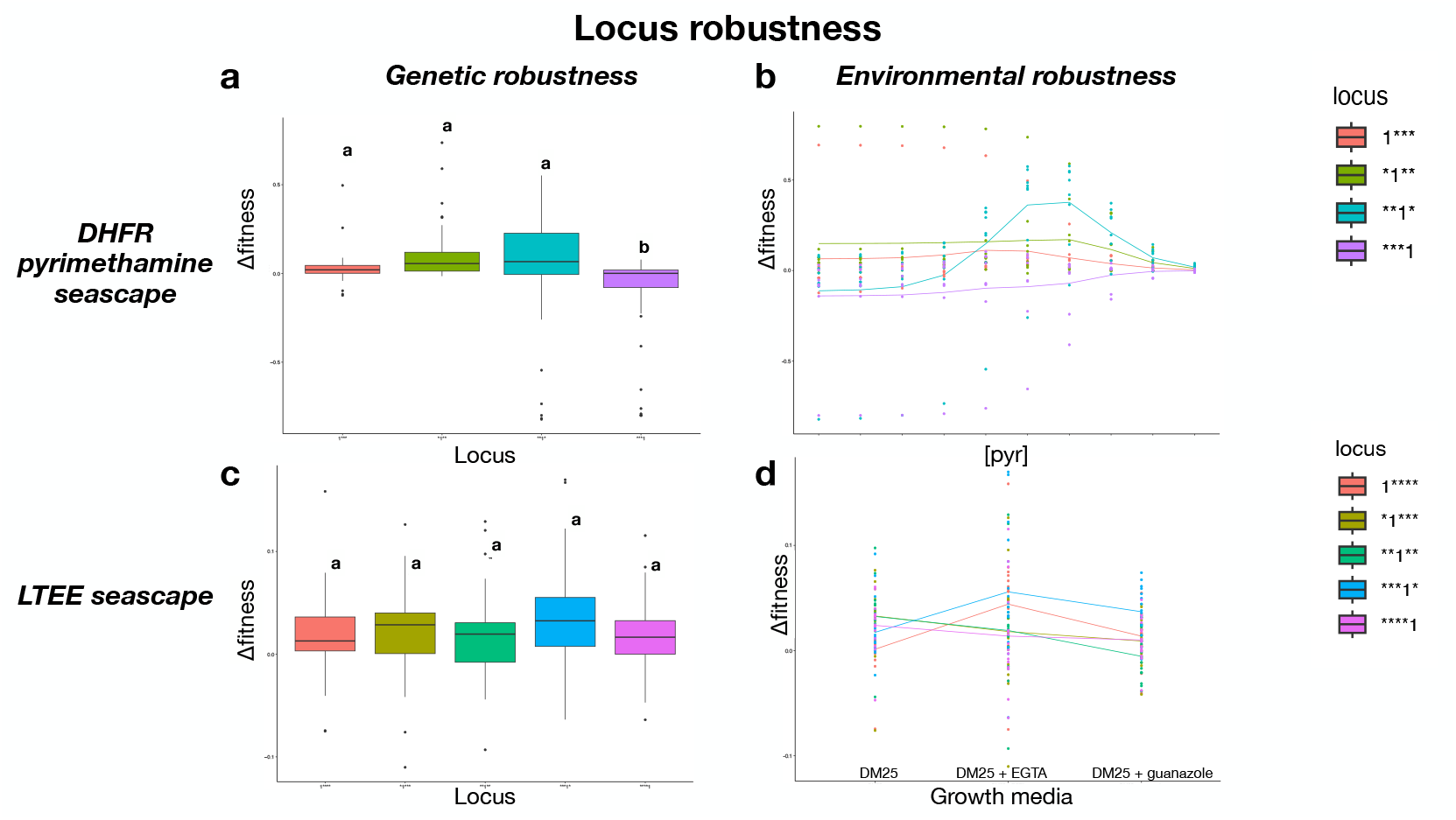
Locus-scale analyses of genetic and environmental robustness for 2^4^ DHFR genotypes in *P. falciparum* across a concentration gradient of pyrimethamine in the DHFR pyrimethamine seascape (top row) and 2^5^ genotypes in *E. coli* across growth conditions in the LTEE seascape (bottom row). (a) and (c) show 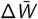 for loci averaged across all genetic backgrounds and environments (genetic robustness) for the DHFR pyrimethamine seascape and LTEE seascape, respectively. (b) and (d) show 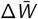 for loci averaged across all genetic backgrounds for each of the individual environmental conditions (environmental robustness) for the DHFR pyrimethamine seascape (for each pyrimethamine concentration) and LTEE seascape (for each of the three growth conditions), respectively.

### 3.3 Properties of interactions between loci

At the scale of locus interactions, the mutation effect reaction norms (MuRNs) demonstrate how mutation effects and epistatic interactions vary across environmental contexts. Across the seascapes, we observe that the MuRNs for individual coefficients (*γ*_*i*_) change in slope across environments, sometimes resulting in MuRNs intersecting (e.g., *γ* **_1_* and *γ*\*_111_ MuRNs intersecting at 10 and 100*µ*g ml^−1^ in the DHFR pyrimethamine seascape), suggesting the presence of G × G × E interactions in the genotype-phenotype map (Figures 4a and c, S3a and c) (*43*). Additionally, in the DHFR pyrimethamine seascape, 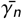 is observed to be highest for the 4th order interaction, followed by 3rd, 2nd, and 1st-order at lower concentrations of pyrimethamine, but at concentrations of 100*µ*g ml^−1^ and above, 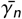 is highest for 1st order interactions, followed by 2nd, 3rd, and then 4th order, with the magnitudes eventually converging (Figure 4b). 5th and 4th order interactions also have a higher 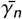 across all environments in the LTEE seascape (Figure 4d). In the *β*-lactam resistance seascape, the magnitude of the coefficient 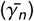 for the 4th order interaction is highest in all environments except for amoxicillin (1024*µ*g ml^−1^)/clavulanic acid (Figure S3b), thus demonstrating how higher-order interactions can shape fitness landscapes across different environmental contexts. In the bla_*TEM*_ cefotaxime resistance seascape, the largest γ_*i*_ is attributed to the single mutation G238S in locus 5, thus mirroring the locus-scale analysis. However, the MuRN also uncovers how the sign and magnitude of pairwise and other higher-order interaction effects fare in comparison to the single locus mutation effects across environments; in particular, in *S. enterica*, the magnitude of the coefficient for the 5th order interaction is higher than 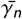 for 1st to 4th order interactions, whereas in *E. coli* and *K. pneumoniae*, the 5th order interaction has the lowest magnitude of all 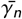 (Figure S3d). All in all, these analyses demonstrate how interactions between loci can differ in sign and magnitude and therefore structure fitness landscapes across different environmental contexts.

**Fig. 4.**
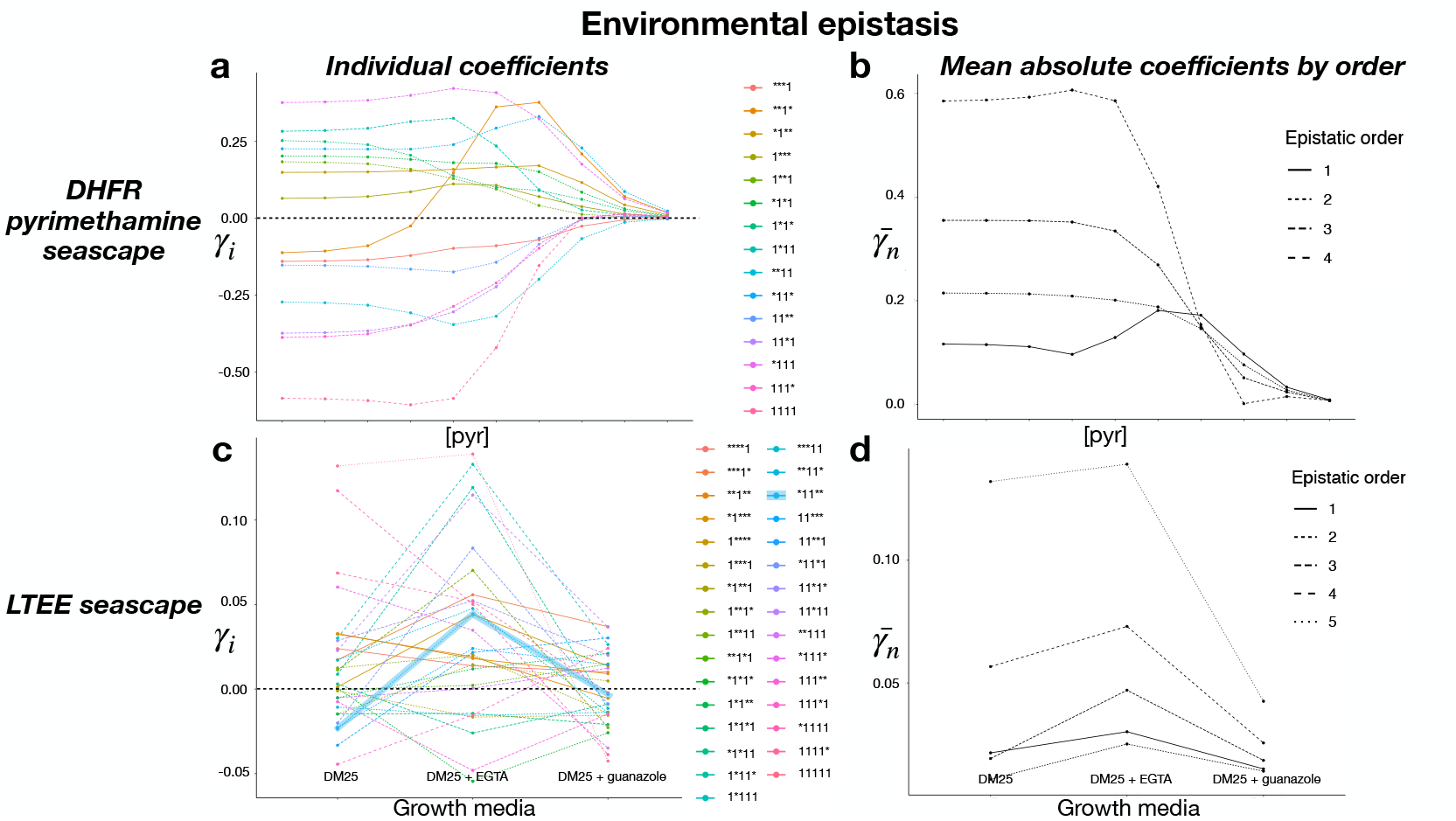
Locus-interaction analyses of environmental epistasis, as computed through the mutation-effect reaction norm (MuRN), for 2^4^ DHFR genotypes in *P. falciparum* across a concentration gradient of pyrimethamine in the DHFR pyrimethamine seascape (top row) and 2^5^ genotypes in *E. coli* across 3 growth conditions in the LTEE seascape (bottom row). Interaction corresponding to each coefficient is defined by the 1s in the binary description (e.g., *11* refers to a 2nd-order interaction between loci 2 and 3, whereas 1*11 refers to a 3rd-order interaction between loci 1, 3 and 4). (a) and (c) show the MuRN depicting coefficients of every possible epistatic interaction (γ_*i*_) for the DHFR pyrimethamine and LTEE seascape respectively (with a black horizontal dashed line to indicate γ_*i*_ = 0), while (b) and (d) show the absolute mean of epistatic coefficients based on epistatic order 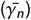 for the DHFR pyrimethamine and LTEE seascape respectively. In (c), the *11** pairwise interaction is highlighted to demonstrate an example of an epistatic interaction changing in sign and magnitude across different environments. Note that for all figures, the 0th-order interaction is not shown.

### 3.4 Properties of alleles

Across seascapes, the *relative* performance of alleles can vary across environmental contexts (Figure 5). Even in seascapes where the rank orders of most alleles remain mostly consistent across environments (e.g., Figure S4a), there are still examples of alleles that change in relative rank order under different environmental contexts, which we discuss in further detail in the next section on landscapes-scale analyses. As well articulated in the literature on reaction norms, the slope and interactions between lines indicate features of the underlying genotype-phenotype map (*69*). For example, crossing reaction norms are representative of (G × E) interactions, which are widespread in both the DHFR pyrimethamine and LTEE fitness seascapes (Figure 5), thus highlighting how the structure of genotype-fitness maps is shaped by interactions between the genotype and the environment.

**Fig. 5.**
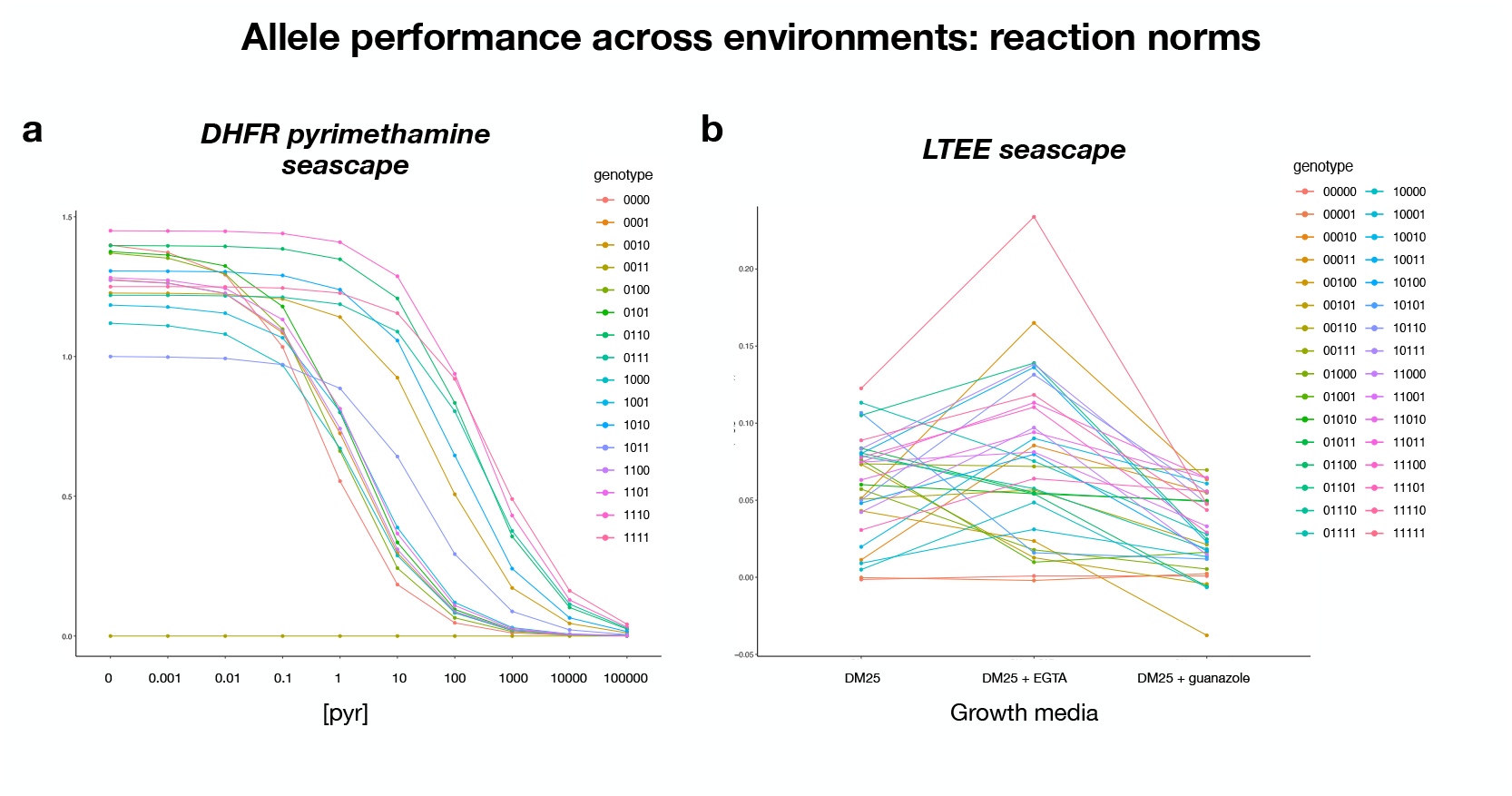
Allele performance across environments: reaction norms. (a) Reaction norms of DHFR pyrimethamine seascape, demonstrating the growth rate of 2^4^ DHFR genotypes in *P. falciparum* across a concentration gradient of pyrimethamine (*43,47*), (b) Reaction norms of LTEE seascape, demonstrate the growth rate of 2^5^ *E. coli* genotypes from the Long-Term Evolution Experiment in 3 different growth conditions (*45,46*).

### 3.5 Properties of trajectories

Doing a qualitative comparison of trajectories across seascapes, we observe that the distributions of *C*_*w*(tot)_ and *C*_*w*(avg)_ differ across seascapes, and within each seascape, different environmental contexts can give rise to different *C*_*w*(tot)_ and *C*_*w*(avg)_ values (Figure 6a, b, d, e; Figure S6a and b). This suggests that the average and/or total within-path competition across all possible evolutionary trajectories can vary depending on the environment, which has broader implications for understanding how adaptation rates across the landscape are dependent on the environment.

**Fig. 6.**
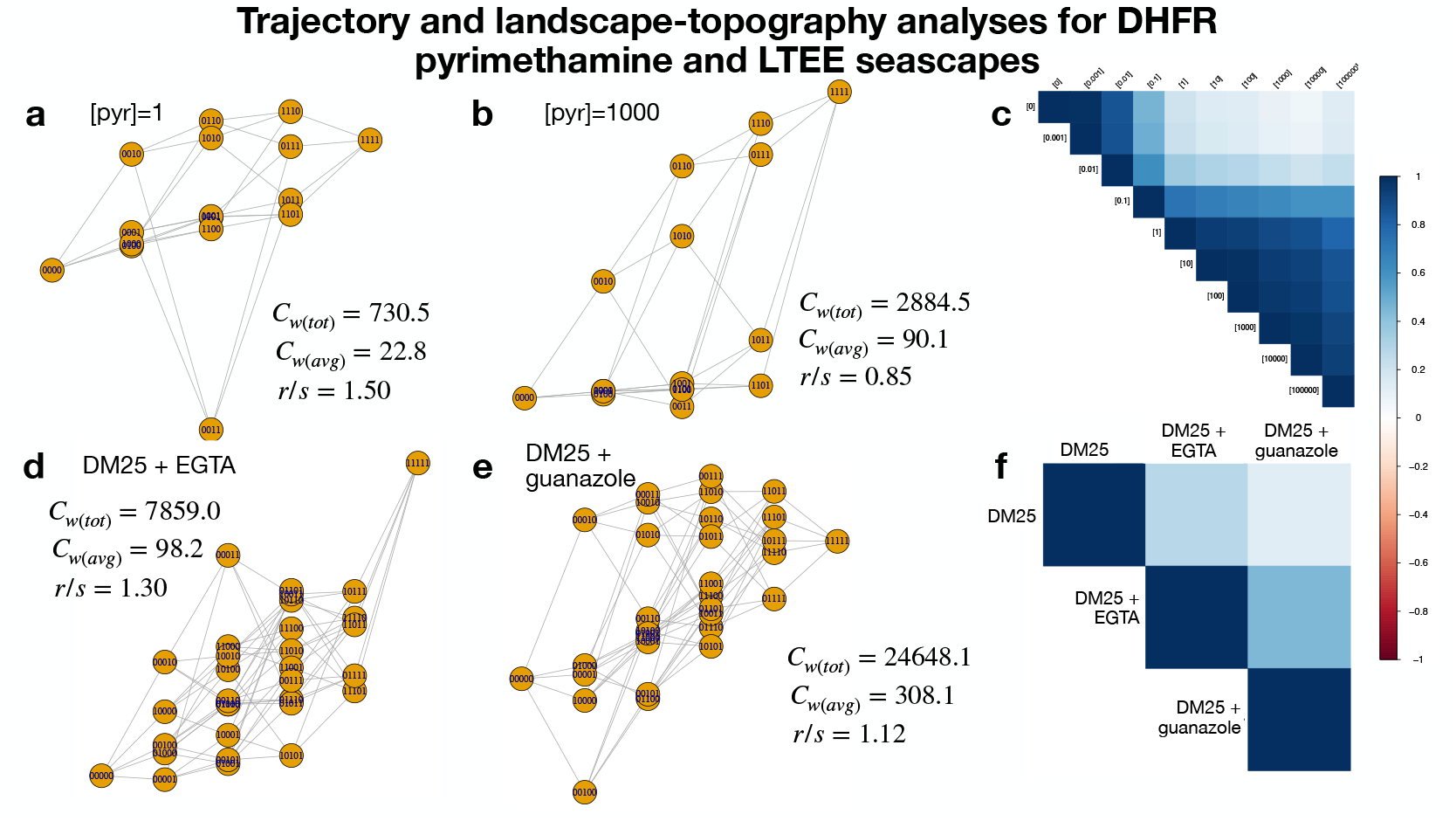
Landscape-scale analyses of evolutionary trajectories and topographies for the DHFR pyrimethamine (top row) and LTEE seascapes (bottom row). (a) Fitness graph with *C*_*w*(avg)_, *C*_*w*(tot)_, and *r* /*s* values for the DHFR pyrimethamine seascape at a pyrimethamine concentration of 1*µ*g ml^−1^. (b) Fitness graph with *C*_*w*(avg)_, *C*_*w*(tot)_, and *r* /*s* values for the DHFR pyrimethamine seascape at pyrimethamine concentration of 1000*µ*g ml^−1^. (c) Kendall’s τ correlation matrix for the DHFR pyrimethamine seascape across the concentration gradient of pyrimethamine. (d) Fitness graph with *C*_*w*(avg)_, *C*_*w*(tot)_, and *r* /*s* values for the LTEE seascape in DM25 + EGTA. (e) Fitness graph with *C*_*w*(avg)_, *C*_*w*(tot)_, and *r* /*s* values for the LTEE seascape in DM25 + guanazole. (f) Kendall’s τ correlation matrix for the LTEE seascape across 3 growth conditions.

### 3.6 Properties of seascape topography

Finally, we qualitatively compare topographical features across the seascapes. When comparing topographical features, we observe that the distributions of roughness-to-slope ratios look different across seascapes, and that within each seascape, environmental contexts can give rise to different extents of ruggedness (Figure 6a, b, d, e; Figure S6c). Furthermore, the Kendall’s τ correlation matrices (capturing ‘seascape’-ness) are different across the 10 seascapes. In some seascapes, the rank orders are highly positively correlated across environments, for example, pyrimethamine concentrations close to each other in magnitude in the DHFR seascape, across the three growth media conditions in the LTEE seascape, and across the three species in the bla_*TEM*_ cefotaxime resistance seascape (Figure 6c and f, Figure S5b). In other contexts, the rank orders are less positively correlated and/or sometimes even negatively correlated between environments such as in the β-lactam resistance seascape (Figure S5a), potentially suggesting trade-off relationships in adapting to different types of beta-lactams and/or different concentrations of beta-lactams (*48,70*).

### 3.7 Other approaches to studying fitness seascapes

Our framework, while covering several scales of granularity through which fitness seascapes can be studied, is not exhaustive. There are many other approaches to studying fitness landscapes that can be applied to fitness landscapes. In this section, we discuss other measures of fitness seascapes that have been proposed and are germane to our central goal of deconstructing fitness seascapes. In this section, we specifically focus on 2 metrics, but we also briefly discuss other metrics in the Supporting Information (see Additional theoretical and empirical approaches for studying fitness seascapes and Table 2). Although these metrics were not formally computed in our study, they can be easily incorporated in future studies. We encourage readers to consider them in their own study of fitness seascapes.

#### 3.7.1 Edge flip fraction

The edge flip fraction is a landscape-scale metric that quantifies the shift in topography of a fitness landscape between two environmental conditions (*71*). Briefly, it calculates the fraction of edge flips between alleles (nodes) for two environments; i.e., how many mutational steps between neighboring genotypes change from beneficial/deleterious to deleterious/beneficial. Computing the edge flip fraction, *ϵ*, involves constructing a transition matrix between nodes for two landscapes, *A* and *B*, to first determine the edge flip matrix, *E* :

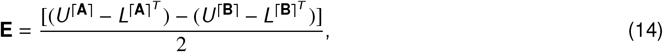

where *U*^⌈**X**⌉^ denotes the upper triangular portion of a matrix **X** with the ceiling function applied to each element, and 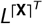 denotes the transposed lower triangular portion of a matrix **X** with the ceiling function applied to each

element. ϵ is then calculated as:

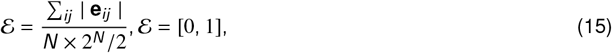

where ϵ = 0 denotes no change in evolutionary direction between two environments and ϵ = 1 denotes change in evolutionary direction for all steps between the environments.

#### 3.7.2 Global epistasis x environment interactions

In our framework, we have discussed how environmental epistasis can alter landscape topography and how we can quantify the contributions of all possible epistatic interactions (pairwise and higher-order). This becomes increasingly complicated to compute in landscapes that are larger than our seascapes, and in landscapes that are not combinatorially complete. Another approach to analyzing how epistasis in adaptive landscapes is modulated by environmental context is to look at *global epistasis*, where the fitness effect of a mutation (Δ*w*) can be predicted by the fitness of the genomic background where it is added (*72*–*74*).

To compute the global epistasis of a given mutation *i*, we can first compute the fitness effect of mutation *i* across the entire set of genetic backgrounds (i.e., the set of genotypes not carrying mutation *i*) for every background *B* in the fitness landscape,

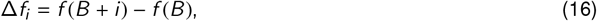

where *f* (*B* + *i*) denotes the fitness of genotype *B* + *i* and *f* (*B*) the fitness of genotype *B* (i.e., genetic background). From the fitness effect, we can then infer different characteristics about the epistasis for a given mutation, such as the directionality of epistasis (slope of linear regression between Δ*f* and *f* (*B*)), the ‘globality’ of epistasis (*R*^2^ of linear regression between Δ*f* and *f* (*B*)), and the strength (magnitude) of epistasis 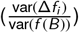. We can therefore adapt our environmental epistasis approach by considering how *global epistasis* is modulated by the environment for larger landscapes, which offers another way of analyzing how interactions between loci are shaped by the environment and therefore, how landscapes change with it. (*75*).

## 4 Discussion

In this study, we examine a set of empirical fitness seascapes with respect to how features vary across environments of various kinds. In doing so, we address an important question: to what extent does the fitness seascape alter our picture of evolutionary genetics? And does the added environmental context offer meaningful information on the dynamics of evolution at the molecular scale? Or is it mostly superfluous? To address this, we compare features of seascapes at multiple scales of granularity: from mutational steps, to mutation effects, and global features of seascape topography. In order to appreciate the importance of the findings, we organize them in order of magnitude of scale, starting with mutational steps, followed by loci, alleles, and landscapes. We find that within individual seascapes and across seascapes, these features change appreciably depending on environmental context, thus highlighting the importance of (a) pivoting from the landscape to the seascape metaphor and (b) studying how these features can vary at different scales of biological organization. In doing so, we advocate for a ‘blueprint’ for studying features of fitness seascapes across different scales of biological organisation, which we summarize in Figure 1 and Table 2.

### 4.1 Implications for evolutionary and population genetics

Our findings demonstrate that environmental context appreciably alters all scales of seascape granularity. At the qualitative level, this might be expected, even obvious. It may therefore be tempting to ask which scale of granularity is most affected by changes across environments. For example, seascapes may shift mutation effects more than features of whole topographies. However, this comparison qualifies as comparing apples to oranges, as each scale of granularity has unique properties. Instead, we address the more concrete question of how environmental context shifts each scale.

In interpreting the specific results of our analysis, several features are notable. The banal observation that allele performance changes across observations is not surprising, and the reaction norm highlights profound amounts of GxE interactions in both the pyrimethamine DHFR and LTEE data sets. This alone demonstrates the importance of the incorporation of environmental information in our descriptions of fitness landscape topography. But a more focused analysis of environmental epistasis and the shape of trajectories highlights the ways that environmental context dictates the pace and shape of adaptive evolution. The mutation-effect reaction norm (MuRN) (Figures 4 and S3) demonstrates the prevalence of GxGxE interactions, where epistatic effects (themselves nonlinear and challenging to predict, by definition) change appreciably across environments (*43*). For example, in the LTEE seascape, the pairwise epistatic interaction between the 2nd and 3rd loci (*11**) has a negative γ_*i*_ in both DM25 and DM25 + guanazole environments, but has a positive γ_*i*_ in the DM25 + EGTA environment (Figure 4c). This demonstrates that epistatic interactions manifest differently in different environments, with implications for the shape of trajectories and landscape topography. Likewise, when comparing how *C*_*w* (avg)_ changes between the DM25 + EGTA and DM25 + guanazole environments (Figure 6d and e), we can infer that populations undergoing adaptive evolution can experience different extents of within-path interference on average, which can have implications for the speed of adaptive evolution across environments (*67*).

Our analysis across scales of granularity is useful because scientists or practitioners might be interested in the fitness seascape metaphor for different questions. For example, those interested in evolvability can focus on individual steps and their features (*36,81,82*). From our results, we can responsibly conclude that the fitness seascape metaphor provides enough variation across context to support its use as a meaningful advance on the fitness landscape, and a useful, (relatively) novel metaphor in population and evolutionary genetics.

### 4.2 Implications for biomedicine and bioengineering

A disproportionate number of studies of empirical fitness landscapes have been conducted in disease systems, specifically those involving antimicrobial resistance (*40,42*–*44,48,51,52,55*). There are multiple reasons for this, including the availability of datasets, which relate to the public health interest in understanding and predicting the evolution of resistance (*83*). The seascape dimension further supports the notion that we can predict, control, or steer adaptive evolution as a means of controlling adaptive evolution. Since drug resistance arises in fluctuating environments, the emergence of the fitness seascape is an essential model for understanding how microbes evolve resistance to small molecules.

Another area where fitness landscapes have been increasingly discussed is bioengineering. The idea here is that the goal of engineering genomes requires knowledge of the underlying fitness landscape, as this can dictate how engineered mutations function with respect to genomic background. The fitness seascape metaphor also aids our perspective. For example, engineering a hypothetical crop with mutations conferring productivity through drought or disease conditions is not a static landscape problem–how the engineered mutations function across environmental contexts is central to whether it will be effective in real-world settings (*84,85*).

Our exploration of practical applications of fitness seascapes suggests a brief reflection on the manner that the technical issues interface with bioethics. Our analysis has revealed how environmental context can shape the fundamental features of the genotype-phenotype map. In some ways, the findings undermine simplistic pictures of genotype-phenotype mapping, the perspective that can yield naive expectations for our ability to control biological organisms. That is, part of the message could be that mutation effects (for example) differ so drastically with even slight environmental variation that we should be thoughtful in our bioengineering pursuits. Such issues have arisen in recent years in discussions of engineering human embryos (*86,87*), as well as in efforts at “de-extinction” (*88*–*92*).

### 4.3 Limitations

Although this study includes a multi-scale analysis of different fitness landscapes, what we have offered does not qualify as a comprehensive treatment of the ways that environmental-dependence can influence features of a fitness landscape. As we have outlined (See Other approaches to studying fitness seascapes section) there are metrics like global epistasis and edge flip fraction that also detail how context-dependence influences evolution at the molecular scale. Nevertheless, we invite others to examine fitness seascapes using whatever approaches might be relevant to their area of interest.

In addition, our study focused on relatively small, combinatorially complete fitness landscapes. Although they are a foundational system in the study of empirical fitness landscapes, new technology has greatly increased the size of fitness landscapes that can be constructed and analyzed (*93*–*99*). Consequently, one may suggest that our proposed framework should be applied to larger data sets. Without digressing into a philosophical debate about the merits of small data versus big data, we offer the subtle claim that early in the rise of a concept (like the fitness seascape), using smaller, well-studied systems can be very useful, as the data have been vetted. Of course, the dichotomy between large and small data is not relegated to questions in evolutionary and population genetics, but is a larger issue that exists across many subfields of biology (and science) (*100,101*).

### 4.4 Conclusions

Our study contributes to a chorus aiming to fortify the fitness seascape as a valuable advance in the fields of evolutionary and population genetics. We provide a mechanistic, multi-scale disentangling of context-dependence and environmental dynamism influencing molecular evolution. This has applications of various kinds, to evolutionary theorists, to clinicians who diagnose and treat diseases, and biological engineers aiming to control how living systems function.

## Acknowledgements

The authors would like to thank members of the OGPlexus, J. Draghi, R. Prum, G. Wagner, and D. Weinreich for helpful conversations on the topic. The authors would also like to thank the organizers and attendees at the 2025 “Molecular Mechanisms of Evolution” Gordon Research Conference, where ideas in this manuscript were discussed.

## Author contributions

SNM and CBO conceived the study, performed the analyses, interpreted the results, and wrote the original version of the manuscript.

## Data and materials availability

All code and data is available at https://github.com/OgPlexus/Seascape1.

## Footnotes

1 It is fair to say that a full examination of every sort of measure of a fitness landscape is far beyond the scope of a single research article; perhaps a future review article can take on this challenge.

2 See referenced manuscript for more details (*67*)

