## Supplementary material for "Deconstructing empirical fitness seascapes across scales of granularity"

#### Contents of this supporting information file:

A Additional theoretical and empirical approaches for studying fitness seascapes ii

B Analysis at the mutational step scale ii

C Analyses at the locus scale for 2 seascapes iii

D Analyses at the locus-interaction scale for 2 seascapes iv

E Analyses at the allele scale for 2 seascapes v

F Analyses at the trajectory and landscape topography scales vi

#### Notes on the Supporting Information section

In this manuscript, we propose a framework for deconstructing the study of fitness seascapes across scales of granularity, primarily focusing on two empirical seascapes. In the supplement, we present analyses of two additional empirical seascapes at different scales of granularity to provide a more comprehensive discussion of the framework. We also provide analyses of all 10 empirical seascapes mentioned in Table 1 at the mutational step and landscape scales to offer general insights on how trends at the mutational step and landscapes-scale have manifested across empirical seascapes.

#### A Additional theoretical and empirical approaches for studying fitness seascapes

Besides the metrics mentioned in the main text of our manuscript, several theoretical and empirical approaches to the study of evolution under fluctuating environments more generally have also been developed. While these approaches do not typically utilize the small, combinatorially complete seascape datasets we used in our framework, they nonetheless address central questions. Some have addressed how the fixation probability of a novel mutation is dependent on the environmental variability a population is subject to (***locus-scale analysis***) (*102*), how mutation rates can evolve in response to environmental and demographic fluctuations (***mutational step, landscape scale-trajectory analyses***) (*103*), how fitness variance of mutations may affect 'environmental memory' and hence evolution across fluctuating environments (***mutational step, locus-scale analyses***) (*104*), and how environmental variability can drive mutational robustness, evolvability, fitness distributions and diversity of populations (***locus, landscape scale-trajectory analyses***) (*105*,*106*).

#### B Analysis at the mutational step scale

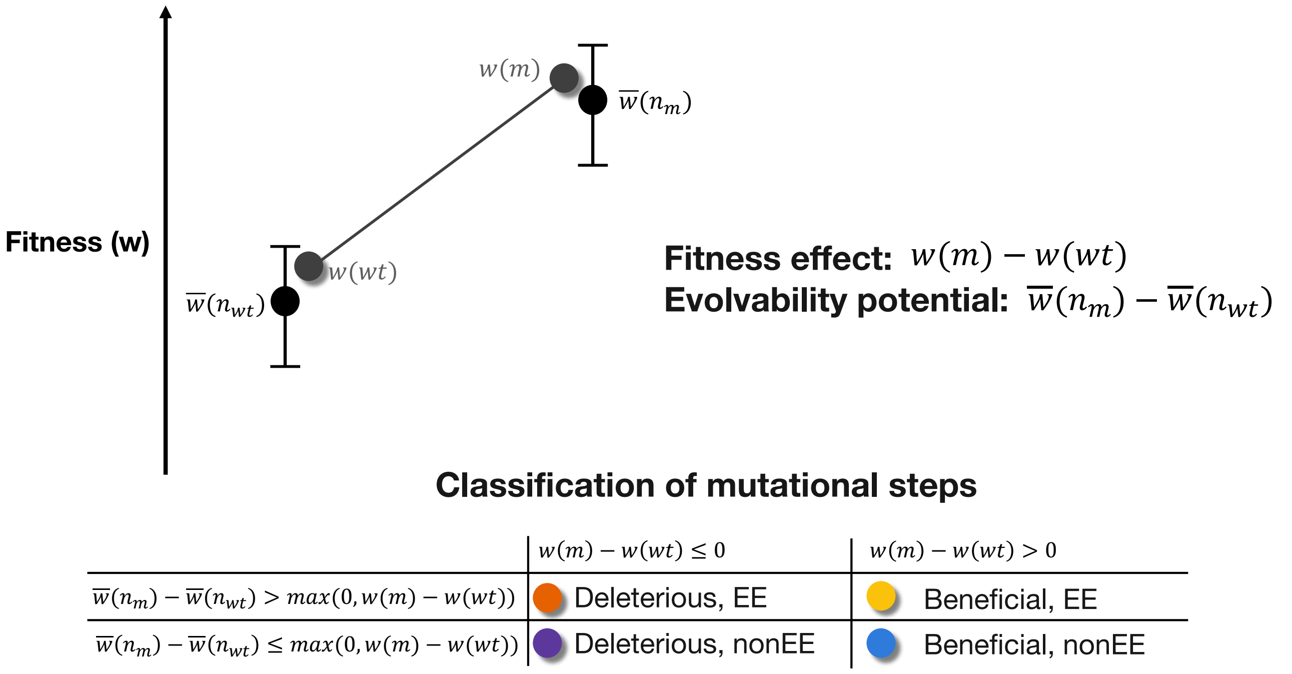


**Fig. S1.** Graphical depiction of how to measure fitness and evolvability at the mutational steps scale, and how mutational steps can be categorized based on their fitness effect and evolvability potential (i.e., whether they are beneficial/deleterious and EE/nonEE). We demonstrate this by taking any given pair of genotypes separated by 1-mutational step and arbitrarily assigning one of the genotypes as the wild-type (wt) and the other as the mutant (m). Figure adapted from Wagner (2023) (*107*).

### C Analyses at the locus scale for 2 seascapes

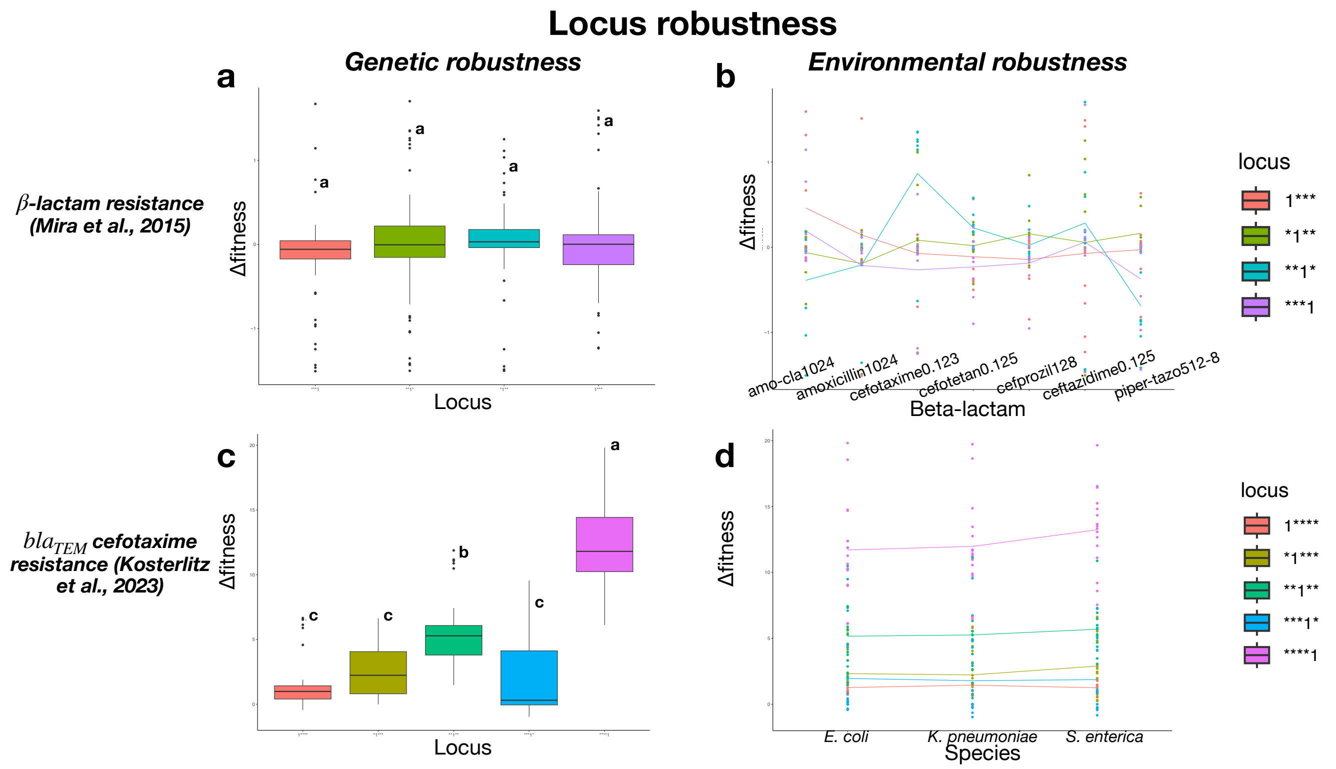


**Fig. S2.** Locus-scale analyses of genetic and environmental robustness for $2^4$ TEM genotypes in *E. coli* across 7 different $\beta$-lactam antibiotics in the $\beta$-lactam resistance seascape (*108*) (top row) and $2^5$ TEM genotypes across 3 bacterial species in the bla$_{TEM}$ cefotaxime resistance seascape (*109*) (bottom row). (a) and (c) show $\Delta\bar{W}$ for loci averaged across all genetic backgrounds and environments (genetic robustness) for the $\beta$-lactam resistance and bla$_{TEM}$ cefotaxime resistance seascapes respectively. (b) and (d) show $\Delta\bar{W}$ for loci averaged across all genetic backgrounds for each of the individual environmental conditions (environmental robustness) for the $\beta$-lactam resistance (each of the 7 $\beta$-lactam antibiotics) and bla$_{TEM}$ cefotaxime resistance (each of the 3 bacterial species) seascapes respectively.

### D Analyses at the locus-interaction scale for 2 seascapes

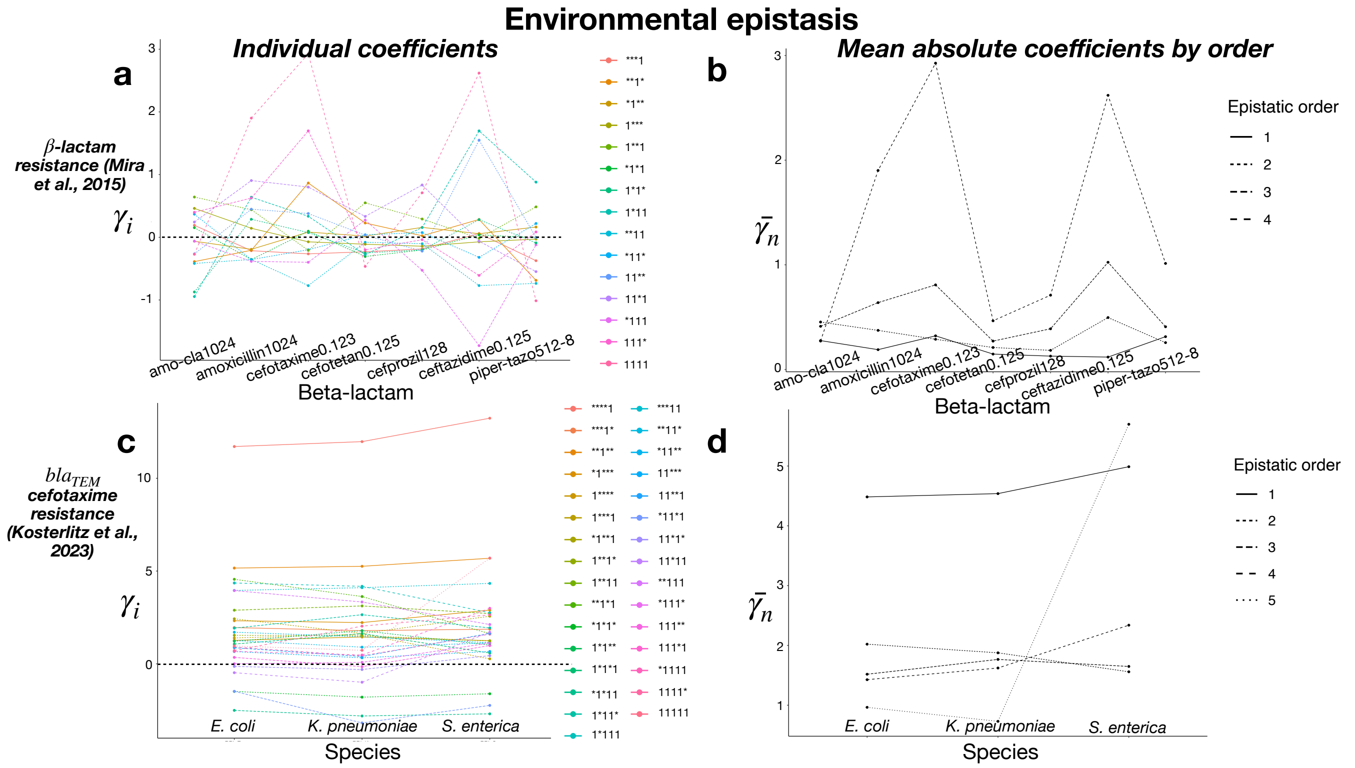


**Fig. S3.** Locus-interaction analyses of environmental epistasis, as computed through the mutation-effect reaction norm (MuRN), for $2^4$ TEM genotypes in *E. coli* across 7 different $\beta$-lactam antibiotics in the $\beta$-lactam resistance seascape (*108*) (top row) and $2^5$ TEM genotypes across 3 bacterial species in the bla$_{TEM}$ cefotaxime resistance seascape (*109*) (bottom row). (a) and (c) show the MuRN depicting coefficients of every possible epistatic interaction ($\gamma_i$) for the $\beta$-lactam resistance and bla$_{TEM}$ cefotaxime resistance seascapes respectively (with a black horizontal dashed line to indicate $\gamma_i = 0$), while (b) and (d) show the absolute mean of epistatic coefficients based on epistatic order ($\bar{\gamma_n}$) the $\beta$-lactam resistance and bla$_{TEM}$ cefotaxime resistance seascapes respectively. Note that for all figures, the 0th-order interaction is not shown.

### E Analyses at the allele scale for 2 seascapes

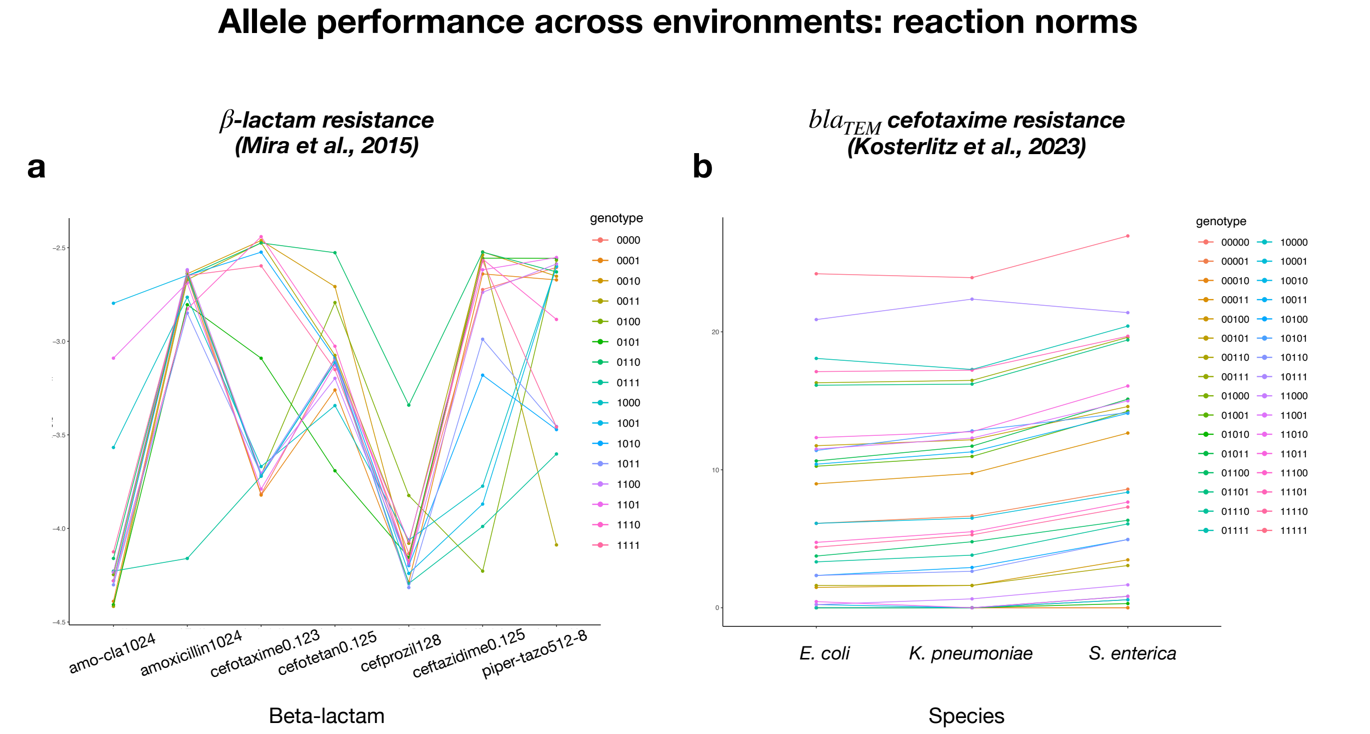


**Fig. S4.** Allele performance across environments: reaction norms. (a) Reaction norms of the $\beta$-lactam resistance seascape (*108*), demonstrating the growth rate of $2^4$ TEM genotypes in *E. coli* across 7 different $\beta$-lactam antibiotics. (b) Reaction norms of the bla$_{TEM}$ cefotaxime resistance seascape (*109*), demonstrating the relative resistance level of $2^5$ TEM genotypes against cefotaxime across 3 bacterial species.

### F Analyses at the trajectory and landscape topography scales

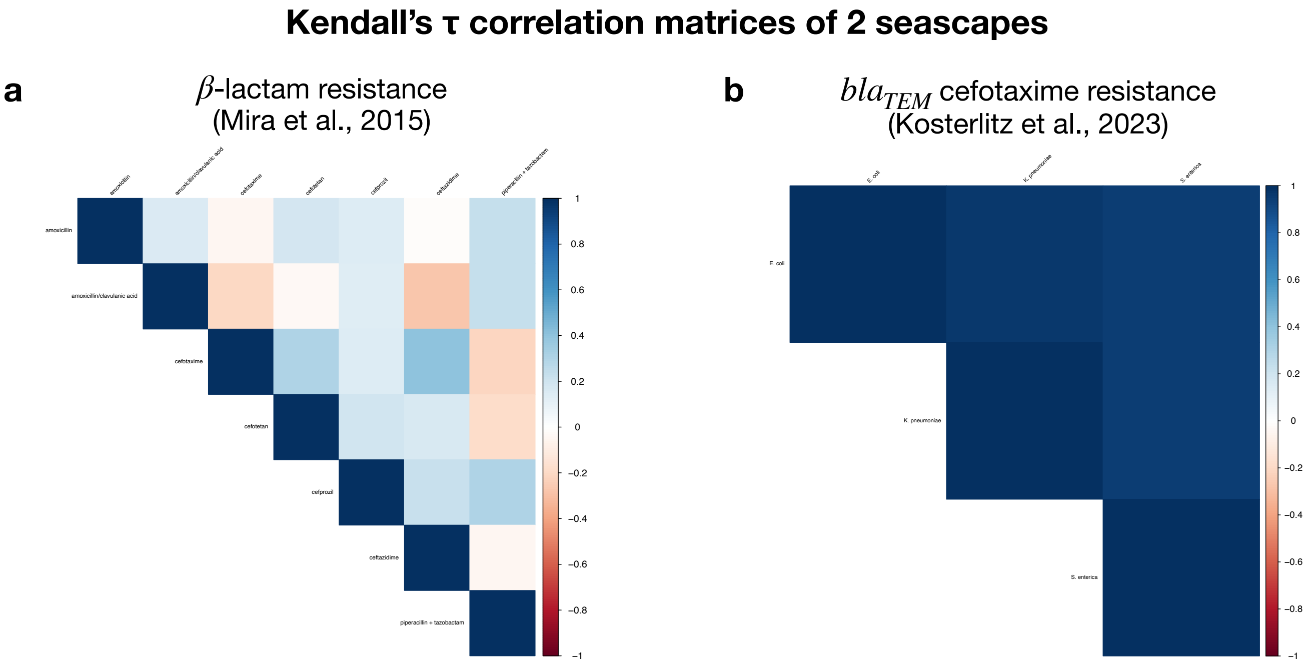


**Fig. S5.** Landscape-scale analyses: Kendall's $\tau$ correlation matrices of genotype rank orders for pairs of environments. (a) Kendall's $\tau$ correlation matrix for 7 different $\beta$-lactam antibiotics in the $\beta$-lactam resistance seascape (*108*), (b) Kendall's $\tau$ correlation matrix for 3 different bacterial species in the bla$_{TEM}$ cefotaxime resistance seascape (*109*).

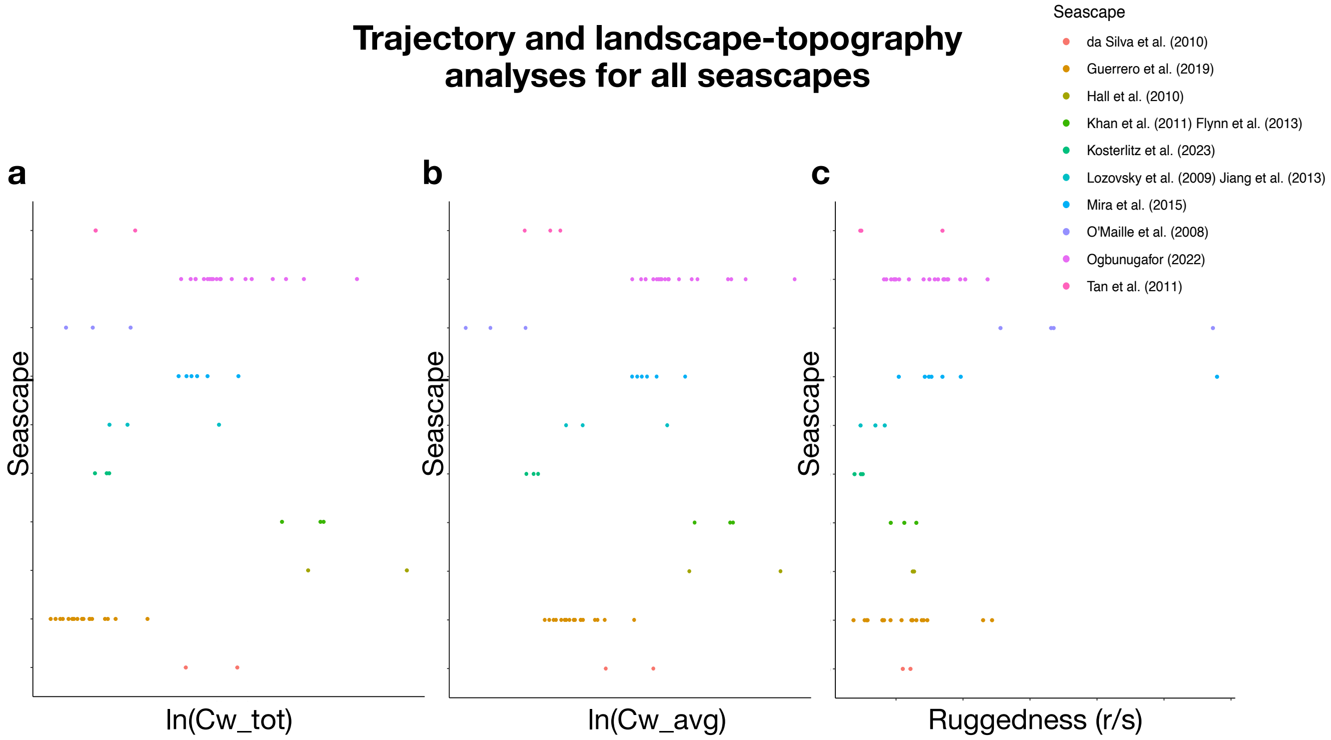


**Fig. S6.** (a) Distribution of $\ln(C_{w(\text{tot})})$ under different environmental and/or phenotypic contexts for all 10 seascapes. (b) Distribution of $\ln(C_{w(\text{avg})})$ under different environmental and/or phenotypic contexts for all 10 seascapes. (c) Distribution of roughness-to-slope ratios under different environmental and/or phenotypic contexts for all 10 seascapes.
